# Chromatin accessibility determines intron retention in a cell type-specific manner

**DOI:** 10.1101/2021.02.17.431609

**Authors:** Veronika Petrova, Renhua Song, DEEP Consortium, Karl J.V. Nordström, Jörn Walter, Justin J.-L. Wong, Nicola J. Armstrong, John E.J. Rasko, Ulf Schmitz

**Affiliations:** Computational BioMedicine Laboratory Centenary Institute, The University of Sydney, Camperdown 2050, Australia; Gene and Stem Cell Therapy Program Centenary Institute, The University of Sydney, Camperdown 2050, Australia; Epigenetics and RNA Biology Program Centenary Institute, The University of Sydney, Camperdown 2050, Australia; Faculty of Medicine and Health, The University of Sydney, Camperdown 2050, Australia; Laboratory of EpiGenetics, Saarland University, Campus A2 4, 66123 Saarbrücken, Germany; Mathematics and Statistics, Curtin University, Bentley, WA 6102, Australia; Cell and Molecular Therapies, Royal Prince Alfred Hospital, Camperdown 2050, Australia

**Keywords:** chromatin accessibility, intron retention, epigenetics, alternative splicing, histone marks, CpG methylation, nucleosome occupancy

## Abstract

Dynamic intron retention (IR) in vertebrate cells is of widespread biological importance. Aberrant IR is associated with numerous human diseases including cancer. Despite consistent reports demonstrating intrinsic sequence features that predispose introns to become retained, conflicting findings about cell type-specific IR regulation demand a systematic analysis in a controlled experimental setting. We integrated matched transcriptomics and epigenetics data (including DNA methylation, nucleosome occupancy, histone modifications) from primary human myeloid and lymphoid cells. Using machine learning we trained two complementary models to determine the role of epigenetic factors in the regulation of IR. We show that increased chromatin accessibility contributes substantially to the retention of introns in a cell-specific manner. We also confirm that intrinsic characteristics of introns are key for them to evade splicing. With mounting reports linking pathogenic alterations to RNA processing, our findings may have profound implications for the design of therapeutic approaches targeting aberrant splicing.

## Introduction

The role of introns in mammalian genomes remains largely unexplained. Given the time and energy required for the transcription and subsequent excision of introns from pre-mRNA, it was important to recognise in recent years that introns can be selectively retained in mature mRNA transcripts and thereby contribute significantly to transcriptomic complexity (Schmitz et al., 2017; Wong et al., 2013). Intron retention (IR) is a form of alternative splicing that was assumed to occur due to the failure of the spliceosome to excise an intron from a pre-mRNA transcript. However, growing evidence suggests that IR is highly regulated by multiple complementary factors (Monteuuis et al., 2019).

IR is widespread across all human tissues and affects more than 80% of protein-coding genes (Middleton et al., 2017). For example, dynamic IR profiles have been identified in key genes involved in hematopoietic cell differentiation and activation (Edwards et al., 2016; Green et al., 2020; Ni et al., 2016; Ullrich and Guigo, 2020; Wong et al., 2013). Fates of intron-retaining transcripts can be diverse and include (i) nonsense-mediated decay triggered by intronic premature termination codons, (ii) detention in the nucleus or nuclear degradation, and (iii) translation into alternative protein isoforms or creation of neoepitopes (Monteuuis et al., 2019; Smart et al., 2018; Wong et al., 2016). A better understanding of how IR is regulated is crucial to determine factors leading to aberrant IR, which has been associated with multiple diseases including cancer (Dvinge et al., 2019; Hershberger et al., 2020; Monteuuis et al., 2020)

Despite numerous studies that describe the role of retained introns in key biological functions in animals and in human diseases (Monteuuis et al., 2020; Monteuuis et al., 2019; Wong et al., 2016), a comprehensive understanding of their regulation is still lacking. Retained introns have conserved intrinsic characteristics such as a higher GC content, shorter lengths, and weaker splice sites in comparison to their non-retained counterparts (Braunschweig et al., 2014; Monteuuis et al., 2019; Schmitz et al., 2017). These features predispose introns to retention but cannot explain the dynamic IR profiles observed in numerous biological processes.

The regulation of alternative splicing has been the focus of many studies. Evidence suggests that alternative splicing is regulated at least at two levels: (i) locally, where *trans*-acting splicing regulators interact with *cis*-acting regulatory elements, and (ii) globally, through the structure of chromatin, which is largely governed by epigenetic factors, including nucleosome assembly, histone modifications and CpG methylation (Zhou et al., 2014).

Previous reports have shown that, apart from intrinsic sequence-based features, intron expression can be regulated through (i) *cis*-regulatory elements, such as sequence motifs attracting *trans*-acting splicing-regulatory RNA binding proteins (Middleton et al., 2017), (ii) core components of the splicing machinery (Wong et al., 2013), and (iii) change in the RNA Pol II elongation rate (Fong et al., 2014). Moreover, an increasing number of studies have found links between epigenetic profiles and IR; reporting that IR is associated with reduced CpG methylation (Gao et al., 2019; Green et al., 2020; Kim et al., 2018; Wong et al., 2017a) and various histone modifications (Guo et al., 2014; Wei et al., 2018). However, these reports have typically established the association of IR with only one epigenetic factor at a time. In general, the question of whether there are dominant epigenetic factors that underpin IR regulation remain unanswered.

In the quest to find a splicing regulatory ‘code’, several studies have used machine learning methods to train models that predict exon usage with increasing precision (Barash et al., 2010; Leung et al., 2014). Moreover, some models were developed to predict cryptic splicing events caused by genetic variations and to link these to human diseases (Baeza-Centurion et al., 2019; Jaganathan et al., 2019; Xiong et al., 2015). However, the computational prediction of IR events has not been attempted to date and the role of epigenetic marks has rarely been considered in computational models of splicing regulation (Monteuuis et al., 2019; Pacini and Koziol, 2018).

In this study, to sought to systematically elucidate the role of epigenetic marks in the regulation of IR. We analysed genome-wide profiles of 6 histone modifications, CpG methylation and nucleosome occupancy at single-base resolution in primary lymphoid and myeloid cells. Using machine learning, we developed two models that predict IR in primary human immune cells. More specifically, we trained a logistic regression with elastic net (EN) classifier and a conditional Random Forest (RF) classifier with matched transcriptomics and epigenomics data from monocytes, macrophages, naïve T-cells, T-central memory, and T-effector memory cells (Figure 1).

**Figure 1.**
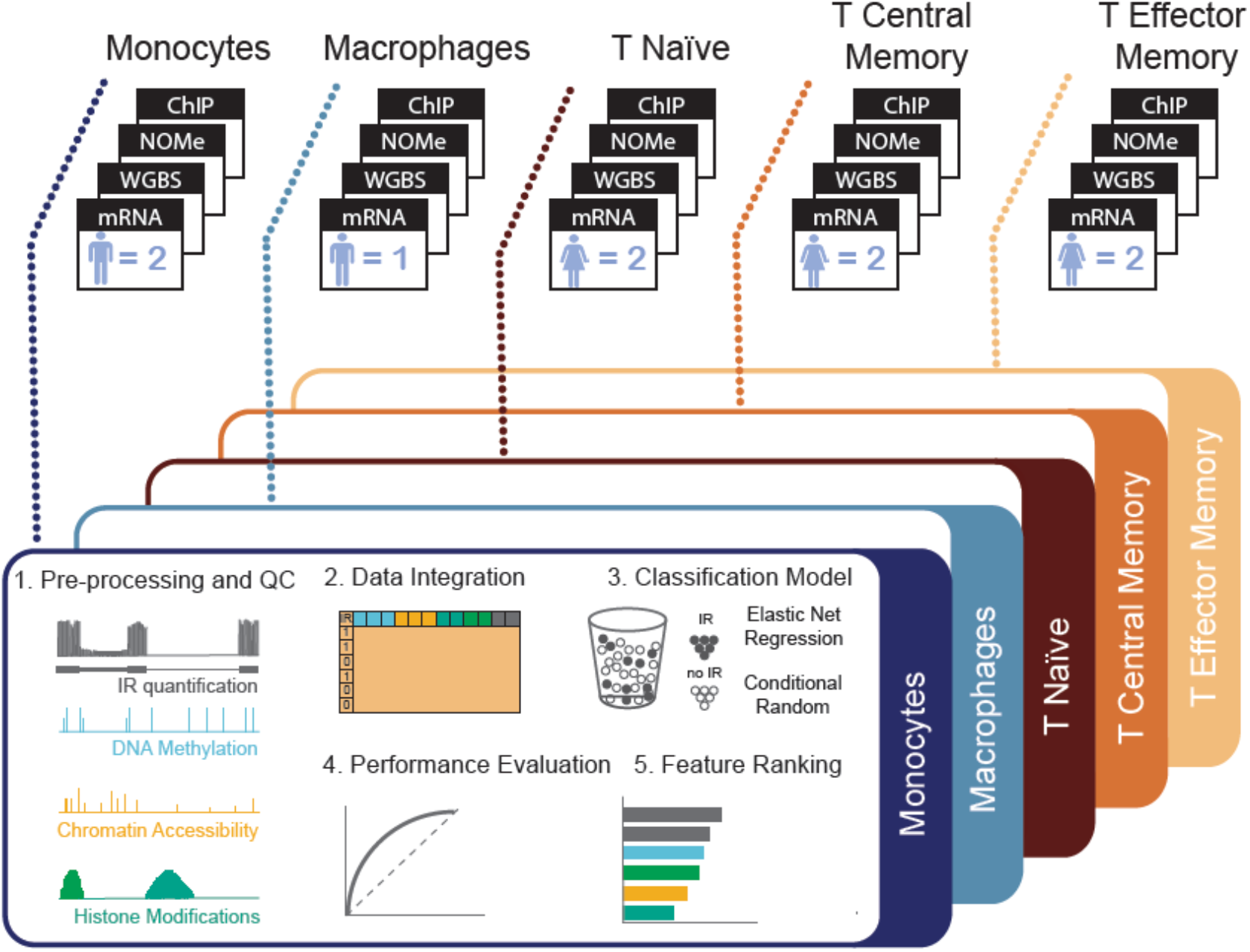
Experimental design and workflow to determine regulators of IR. Raw high-throughput data were processed for each biological replicate and amalgamated by cell type from the indicated number of samples (n). The output was used for feature extraction: IR events were treated as a binary outcome and we trained an Elastic Net (EN) regression model and a conditional Random Forest model with a total of 46 sequence-based and epigenetic features. Using feature ranking, we identified the factors that were most strongly associated with IR outcomes and compared the performances of both modelling strategies. These steps were repeated for each cell type.

Our results show that intrinsic characteristics are key for introns to evade splicing and that epigenetic marks may modulate IR levels in a cell type-specific manner, where the dominant factor for dynamic IR regulation is chromatin organisation.

## Results

### Intrinsic features of retained introns are consistent across cell types

To investigate how IR is regulated in primary immune cells (CD4+ T-cells, monocytes, and macrophages), we integrated transcriptomics (mRNA-Seq) data with epigenomics data including genome-wide CpG methylation (WGBS), histone modifications (ChIP-Seq), and nucleosome occupancy (NOMe-Seq) (Table S1). The cells were isolated from peripheral blood of 2 healthy donors, except for the monocyte-derived macrophages. Using the IR identification software IRFinder (Middleton et al., 2017), we quantified IR events of expressed genes (FPKM>1) in five cell types across myeloid and lymphoid cells, representing two modes of differentiation: monocyte-to-macrophage differentiation and naïve T-cell differentiation into central memory (CM) and effector memory (EM) T-cells. Introns that were present in at least 10% of a gene’s mature mRNA transcripts (IR_ratio_ ≥ 0.1) with an overall intron depth ≥ 10 were considered retained. Non-retained introns were defined as those with an IR_ratio_ ≤ 0.01 and intron depth < 10.

We identified a total of 26,147 retained introns in 12,379 genes, some of which were retained in both myeloid and lymphoid cells while others were cell type-specific (Figure S1A). Consistent with previous reports, retained introns in our dataset are shorter in length, exhibit a higher GC content and weaker splice site strengths compared to non-retained introns (Figure S1B-E). Our analysis revealed diverse splicing patterns in myeloid and lymphoid cells. While 40% of the retained introns in myeloid cells were significantly differentially retained (ΔIR ≥ 0.1; p < 0.05 Audic-Claverie test) between monocytes and macrophages (571/1425), T cells displayed greater stability in regard to IR with only 8% of introns classified as differentially retained (146/1812 in naïve T vs CM, and 80/969 in CM vs EM). In contrast to the monocyte-to-macrophage differentiation, where we observed a reduction in IR events (Figure 2A), the overall number of retained introns remained consistent in all CD4+ T cells. These patterns coincide with fewer changes in gene expression during T cell differentiation in contrast to major gene expression changes in monocyte-to-macrophage differentiation (Figure S1F).

**Figure 2.**
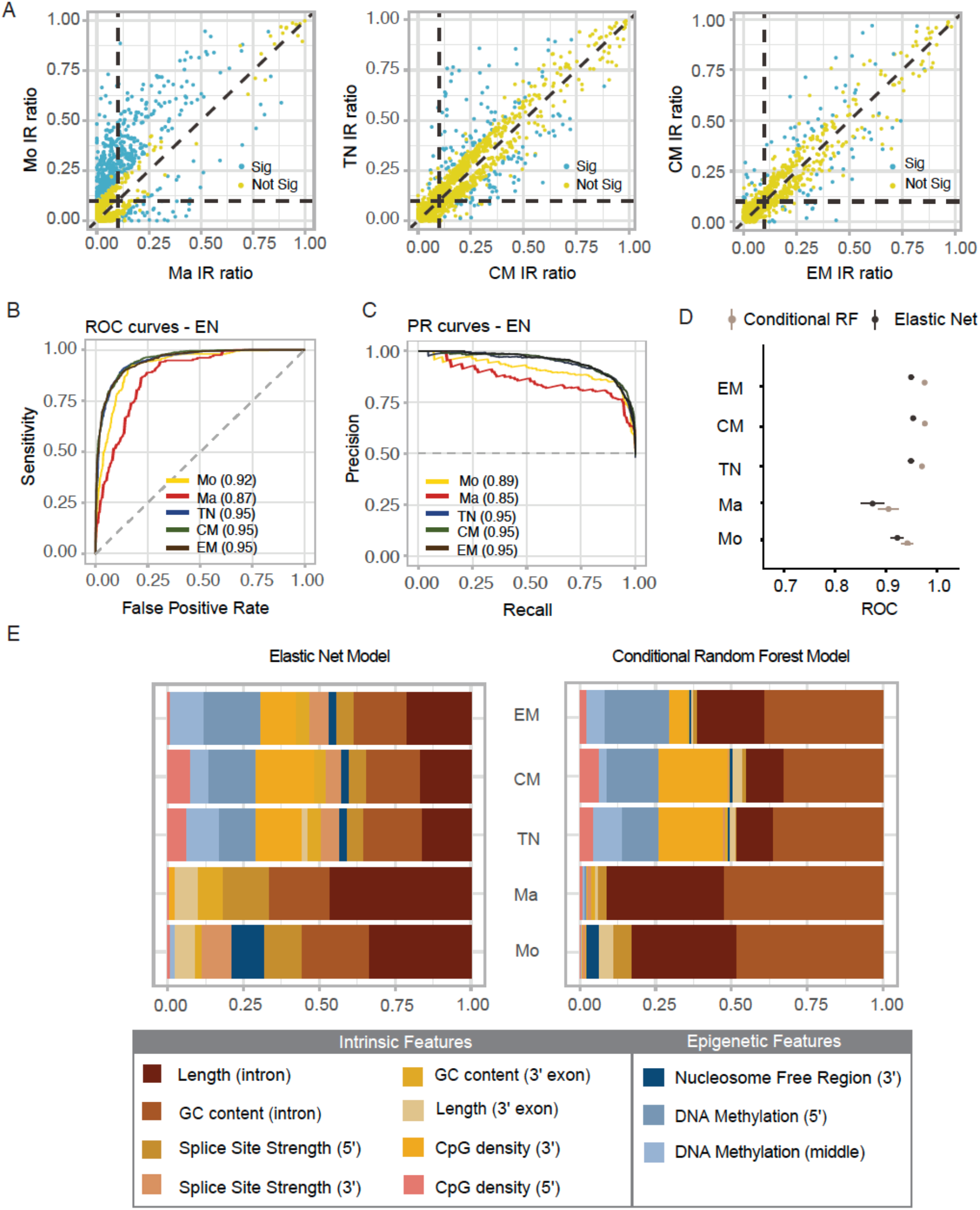
IR prediction and model feature association analyses. **(A)** Scatter plot of differential IR events (Sig blue – significant; Not Sig yellow – not significant) between monocytes (Mo) vs macrophages (Ma) (left), Naïve (TN) vs Central Memory (CM) T cells (middle), and Central Memory vs Effector Memory (EM) T cells (right). **(B)** Receiver operating characteristic (ROC) curves and **(C)** precision recall (PR) curves comparing the performance of the EN classifier in five cell types. **(D)** Comparison of AUC values between EN and cRF algorithms, error bars show 95% confidence interval. **(E)** Variable importance scores for the top 10 features identified by EN and conditional RF algorithms. The scores were scaled to values that add up to 1.0 and the size of a bar corresponds to the effect size.

Most retained introns in our analysis overlapped with histone marks (HM) or with a nucleosome free region (NFR, predicted from NOMe-seq data) around their 5’ and 3’ splice sites (+/100 bp) as well as the middle of an intron (Figure S2A). Interestingly, many non-retained introns (~50%) lacked such epigenetic marks in lymphoid cells (as opposed to only 20-30% of retained introns). H3K36me3 was the most frequently observed histone modification followed by NFR peaks. In retained introns, between 30% and 60% of H3K36me3 signals were classified as strong (see Methods), whilst in non-retained introns the proportion of overlap with the regions of strong signal ranged between 2% and 18%. Again, the patterns of signal strength varied between the cell types (Table S3).

CpG methylation profiles (extracted from WGBS data) for retained and non-retained introns displayed a characteristic bimodal distribution with two distinct peaks at 0% and 100%. Differential methylation was predominantly found at the splice sites when we compared regions of genomic DNA associated with IR and no IR. At the 5’ splice sites, we observed higher methylation levels in retained compared to non-retained introns in all five cell types. However, this trend was reversed in the lymphoid cells at the 3’ splice sites and in the middle of introns (Figure S2B).

M.CviPI enzyme, used in NOMe-seq experiment, methylates cytosine dyads in GC sequence and GCH methylation levels (where H is any nucleobase except guanine) provide information about chromatin accessibility. Unlike endogenous CpG methylation, GC dinucleotides are rarely fully methylated, therefore the mid-range levels (anywhere between 20 to 50%) are usually sufficient to indicate open chromatin regions. In our data, chromatin accessibility (i.e. GCH methylation) increased from monocytes to macrophages with slightly higher levels in retained introns, while lymphoid cells had increased chromatin accessibility (GCH methylation levels 15-35%) but with lower levels in retained introns compared to non-retained introns (Figure S2C).

To determine important factors of IR regulation, we compiled sequence-based and epigenetic features: (i) sequence-based features: intron length, GC content, splice site strength, CpG density (also referred to as intrinsic features), (ii) transcriptomics features: percent spliced-in (PSI) values of the flanking exons, and (iii) epigenomics features extracted from the WGBS, ChIP-Seq (H3K9me3, H3K27me3, H3K27ac, H3K36me3, H3K4me1, H3K4me3), and NOMe-Seq data (Table S2). We then used these features (n=46) to train EN models for each cell type and predict whether introns are either retained or non-retained. The performance of our models was assessed based on the area under the receiver operating characteristic curve (AUC) values, which ranged between 0.87 and 0.95 (Figure 2B) and values for the area under the prediction-recall curve (accuracy) ranging between 0.85 and 0.95 (Figure 2C). The consistently high values suggest that the model choice was appropriate for the task.

Next, in order to evaluate whether the learned relationship between the model features and IR was generalizable across cell types we trained our model with data from one cell type and tested it with data from another cell type. For all training/test data pairs, the AUC and accuracy metrics were comparable to those models that were trained and tested on the same cell type (Table S4).

The EN model assumes a monotonic linear relationship between the class variable and the model features. To determine whether this assumption is adequate for IR classification, we also trained conditional random forest (cRF) models, which do not make any prior assumption about the relationship between the outcome of interest and the model features. Comparing the results from both types of models, we found that cRF performed slightly better than EN with AUC values ranging between 0.91 and 0.98 (Figures 2D, S3A) and PR values between 0.87 and 0.95 (Figure S3B).

To assess which features contribute most to the model performance (and thus, the relevance of a feature to IR), we used variable-importance measures (VIM). For EN, these are the regression coefficients ordered from lowest to highest, where parameters with larger values have a greater effect. For cRF, variable importance was calculated as the mean decrease in accuracy after permutation of each model feature (Figure 2E). Given the known properties of retained introns it was no surprise that intrinsic features, such as length, GC content and CpG density were ranked as the top predictors with a high level of agreement across all cell types analysed. Again, we observed consistency between the EN and cRF models, except for minor variations in the order that important features were ranked in.

Epigenetic features were also ranked among the top 5 predictors across all models and cell types, however their nature and relative importance varied between cell types (Figure 2E). Overall EN models ranked epigenetic features as moderately to very important (VIM between 0.4 and 0.8), which is comparable to the intrinsic features (ranging between 0.3 and 1). In contrast, cRF identified epigenetic features as somewhat important with VIM mostly below 0.50 (Figures S3C, S3D). Nevertheless, intrinsic features were consistently identified as most relevant for correctly classifying IR, suggesting that these features predispose introns to being retained irrespective of cell or tissue type.

### Chromatin accessibility is predicted to be the strongest regulator of IR

In the previous section we classified IR on a cell type-specific basis and determined the intrinsic features as having the strongest association with IR outcomes. However, we often find that an intron is retained in one cell type but not in another. In those cases, factors beyond intrinsic features are the likely drivers of this transition.

To find these IR determinants, we modified our initial modelling approach by focusing only on the dynamic introns - those that changed their retention status between cell types (Figure 3A). In total, 1,540 introns matched this criterion with various IR patterns (Figure 3B). We used these introns to train EN and cRF models with both epigenetic and intrinsic features. The cRF model was performed superior to the EN model achieving AUCs of 0.85 and 0.76, respectively (Figure 3C). cRF also achieved a higher area under the precision-recall curve value (0.83) than EN (0.73) (Figure 3D). The poorer performance of EN might be a reflection of the model’s inability to fully utilise complex structures within the omics data, thus supporting the notion that a relationship between chromatin modifiers and IR is indeed nonlinear, as previously suggested (Singer et al., 2015).

**Figure 3.**
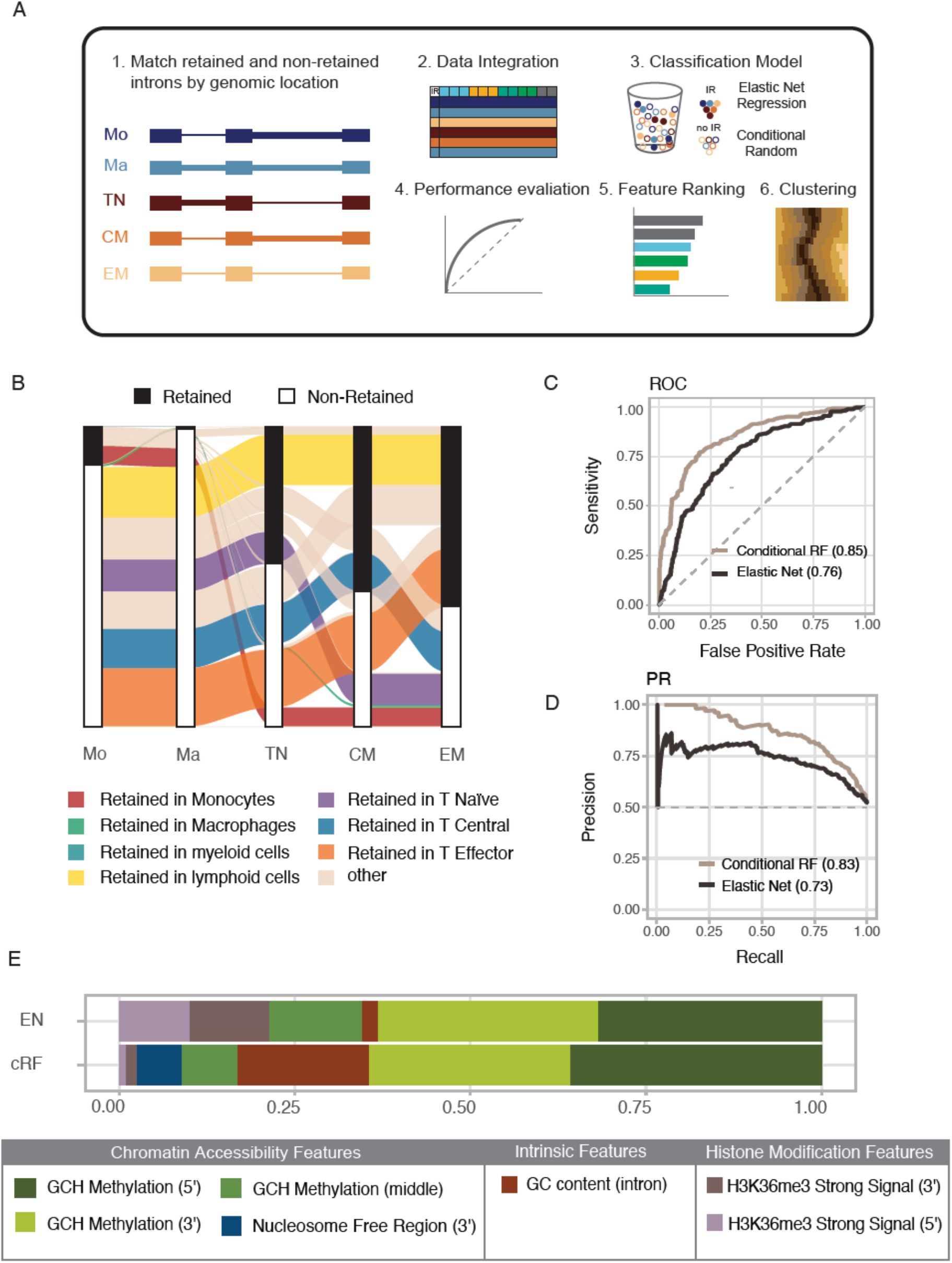
Analysis of dynamics intron retention. **(A)** Modified modelling strategy from Figure 1. Only introns that were found to be in retained and non-retained states in different cell types were included in the analysis. **(B)** Alluvial plot illustrating the dynamics of IR states among the five cell types. **(C)** ROC and **(D)** PR curves comparing the performance of cRF (brown) and EN (black). **(E)** Variable importance scores for the top 5 features identified by EN and conditional RF algorithms, scaled between 0 and 1.

Evaluation of feature rankings revealed that, despite varying model performances, both EN and cRF models identified features related to chromatin accessibility as most important for correct IR classification (Figure 3E). These features include GCH methylation and GCH (i.e. nucleosome) occupancy and the presence of nucleosome free regions (NFRs). GCH methylation at the 5’ and 3’ splice sites were determined as most important features discriminating retained form non-retained introns in both models. The cRF classifier also identified CpG methylation as somewhat important for IR classification, which has a known relationship with chromatin accessibility (Farlik et al., 2016; Lay et al., 2015; Taberlay et al., 2014). Interestingly, the cRF model also identified GC content as a moderately important contributor to IR outcomes, whilst the EN model included histone marks (H3K27ac and H3K36me3) in their top 10 predictors (Figure S4A).

### Epigenetic IR regulation is independent of gene expression regulation

It is reasonable to assume that changes in the epigenomic landscape might not directly affect IR but rather gene expression (Jaenisch and Bird, 2003). To confirm that the features identified as relevant to IR are independent from gene expression regulation, we split dynamically retained introns into three groups: (i) host gene expression is reduced along with the change in IR status, (ii) host gene expression remained stable (log_2_ FC FPKM ≤ 2), and (iii) host gene expression increased (Figure 4A). For most of the dynamic introns the host gene expression remained unchanged (N = 1,220), whilst down- and upregulated host genes were associated with 73 and 247 alternately retained introns, respectively. We repeated the classification analysis on the group of introns where the IR changes were not accompanied by host gene expression changes. Since the relationship between IR and epigenetic model features is not linear, as was established in the previous section, we only used the cRF algorithm.

**Figure 4.**
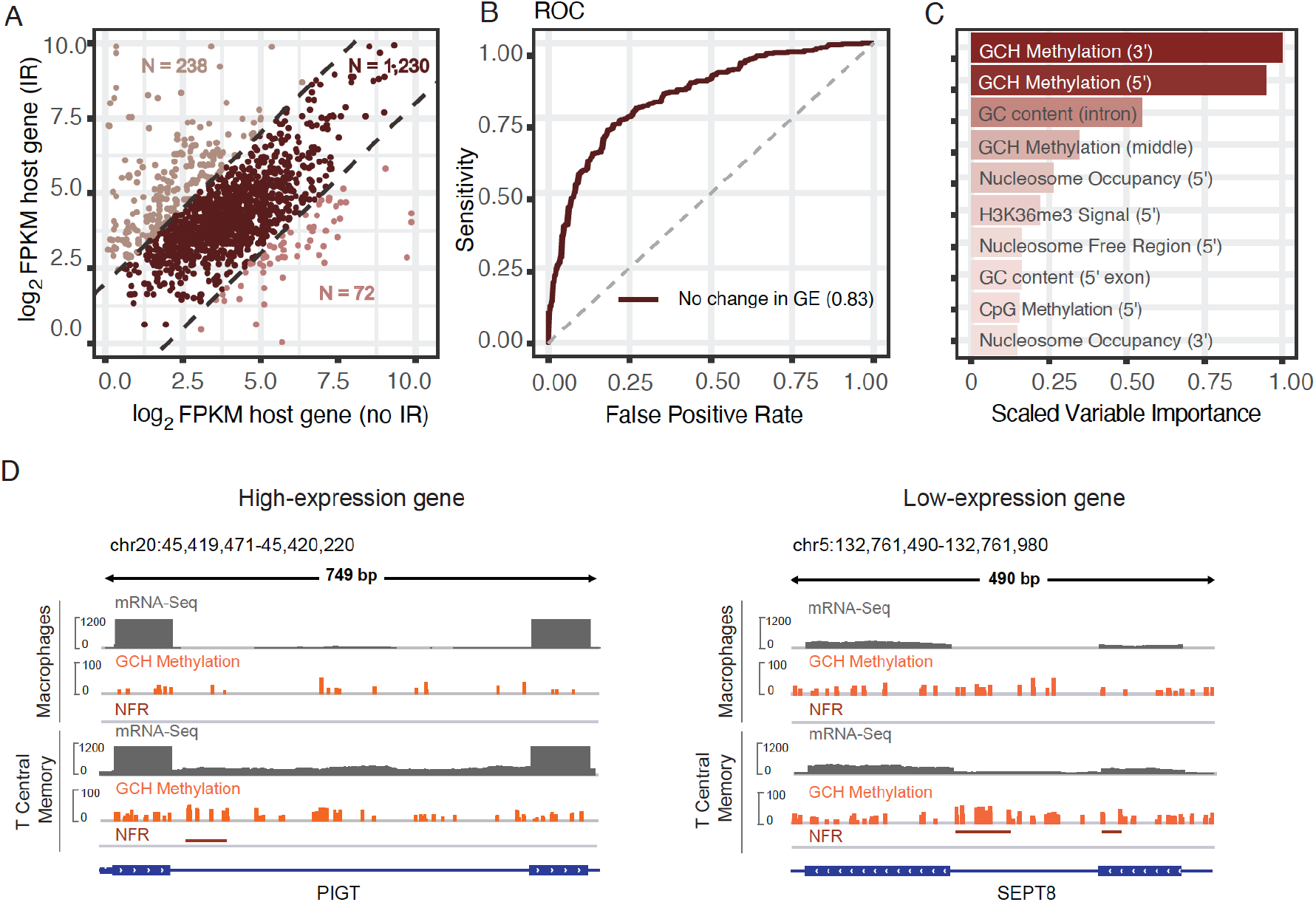
Analysis of introns from genes with non-differential expression levels. **(A)** Scatter plot of host gene expression for introns that change their IR status. **(B)** ROC curve indicating the performance of a conditional RF model fitted on the data from non-differentially expressed genes (GE, gene expression). **(C)** Ranking of the features based on the scaled variable importance scores. **(D)** Integrative Genomics Viewer (IGV) plots revealing higher density and hypermethylation levels of GCH sites in the splice site regions of differentially retained introns in both highly- and lowly-expressed gene examples (NFR – Nucleosome Free Region, GCH Methylation – methylation levels of GC dinucleotides followed by any nucleobase except guanine).

The model fitted on this data subset achieved an AUC of 0.83 (Figure 4B) and an area under the precision-recall curve value of 0.78 (Figure S4B). The features that were selected as important were GCH methylation at the 5’ and 3’ splice sites and GC content in the same order as in the model trained on all dynamically retained introns (Figure 4C). This observation held true for both highly and lowly expressed host genes (Figures S4C). We therefore concluded that the observed epigenetic changes associated with IR modulation are independent from gene expression regulation. In Figure 4D, we show two exemplary introns where greater chromatin accessibility was associated with an increase in IR: Phosphatidylinositol Glycan Anchor Biosynthesis Class T (PIGT) helps building the glycosylphosphatidylinositol-anchor which is found on the surface of various blood cells (Figure 4D, left). PIGT is known to express many isoforms through alternative splicing including IR. The nucleotide binding protein SEPTIN8 is a regulator of cytoskeletal organization, which has multiple alternatively spliced transcript variants as well (Figure 4D, right).

### Dynamic changes in chromatin structure are responsible for cell type-specific IR

As chromatin accessibility was identified as the strongest predictive factor for differential IR, we closely examined its relationship with retained and non-retained introns. We identified 5 distinct GCH methylation profiles in the +/- 200 bp region around the 5’ splice site of retained introns (Figure 5A, left). Similar clustering profiles were identified in the region around 3’ splice sites and the middle of introns (Figure S5). To understand changes in chromatin status in the context of differential IR, we plotted the GCH methylation values of the same introns when they were not retained (Figure 5A). The associated heatmap shows that GCH methylation is widely depleted in non-retained introns, with no distinct clustering. In retained intron, however, we observed a clear increase in GCH methylation immediately upstream or downstream from the 5’ splice site (Figure 5B, clusters 1, 3 and 4). We also identified a group of retained introns with relatively low levels of GCH methylation (cluster 2) and another with particularly strong GCH methylation (cluster 5).

**Figure 5.**
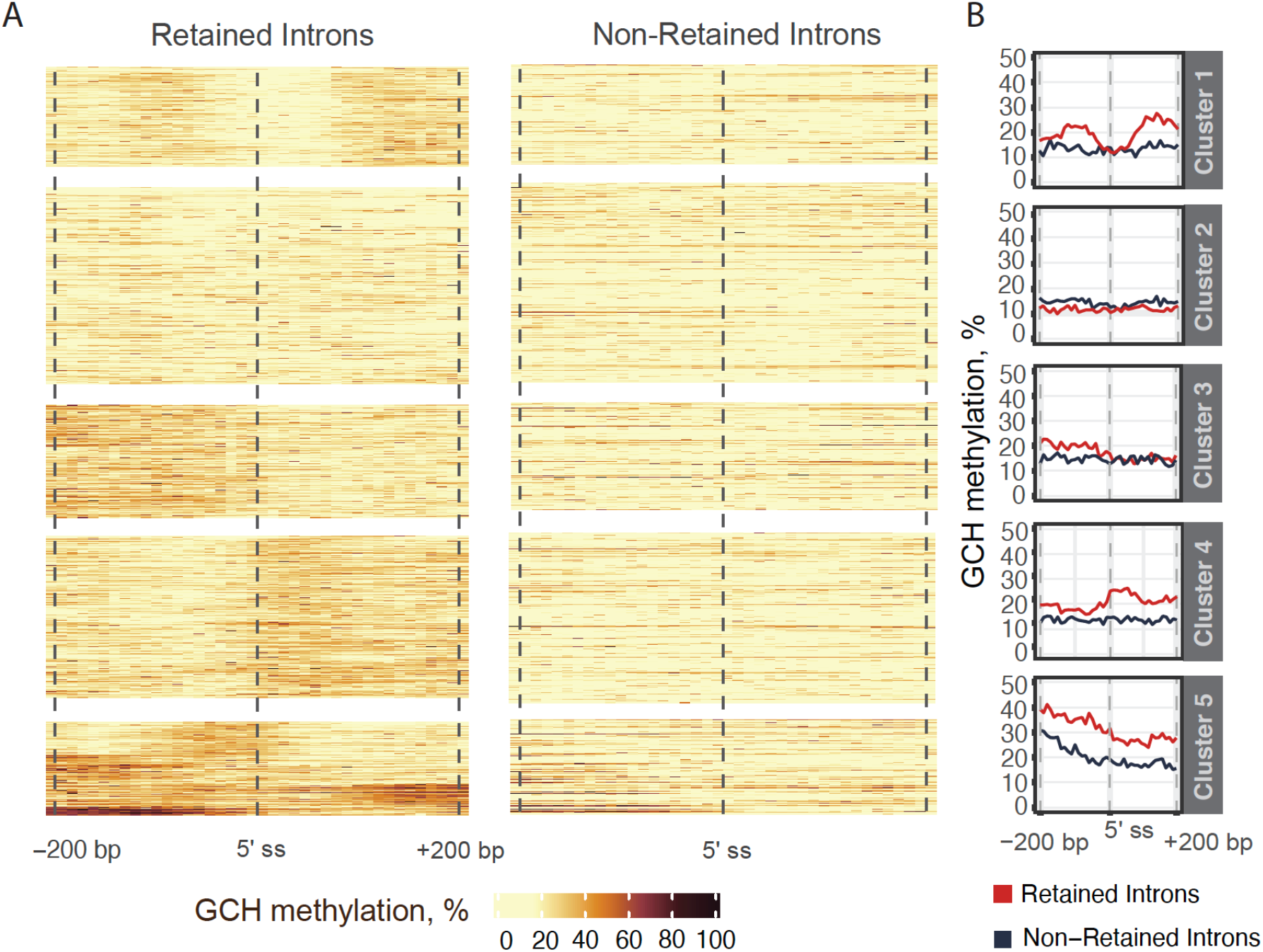
GCH methylation clustering in differentially retained introns. **(A)** Clustering of GCH methylation in the +/- 200 bp region around the 5’ splice site (ss). Each line corresponds to one intron that is either in a retained (left) or non-retained state (right). **(B)** Line plots showing average GCH methylation values (i.e. chromatin accessibility) in retained vs non-retained introns across 5 clusters.

Upon visualising the intronic regions that changed their IR status between cell types, we observed greater chromatin accessibility levels in retained introns (Figure 6A). Moreover, for the majority of introns, we found that IR gain was accompanied with a reduction in H3K36me3 signal (Figure 6A).

**Figure 6.**
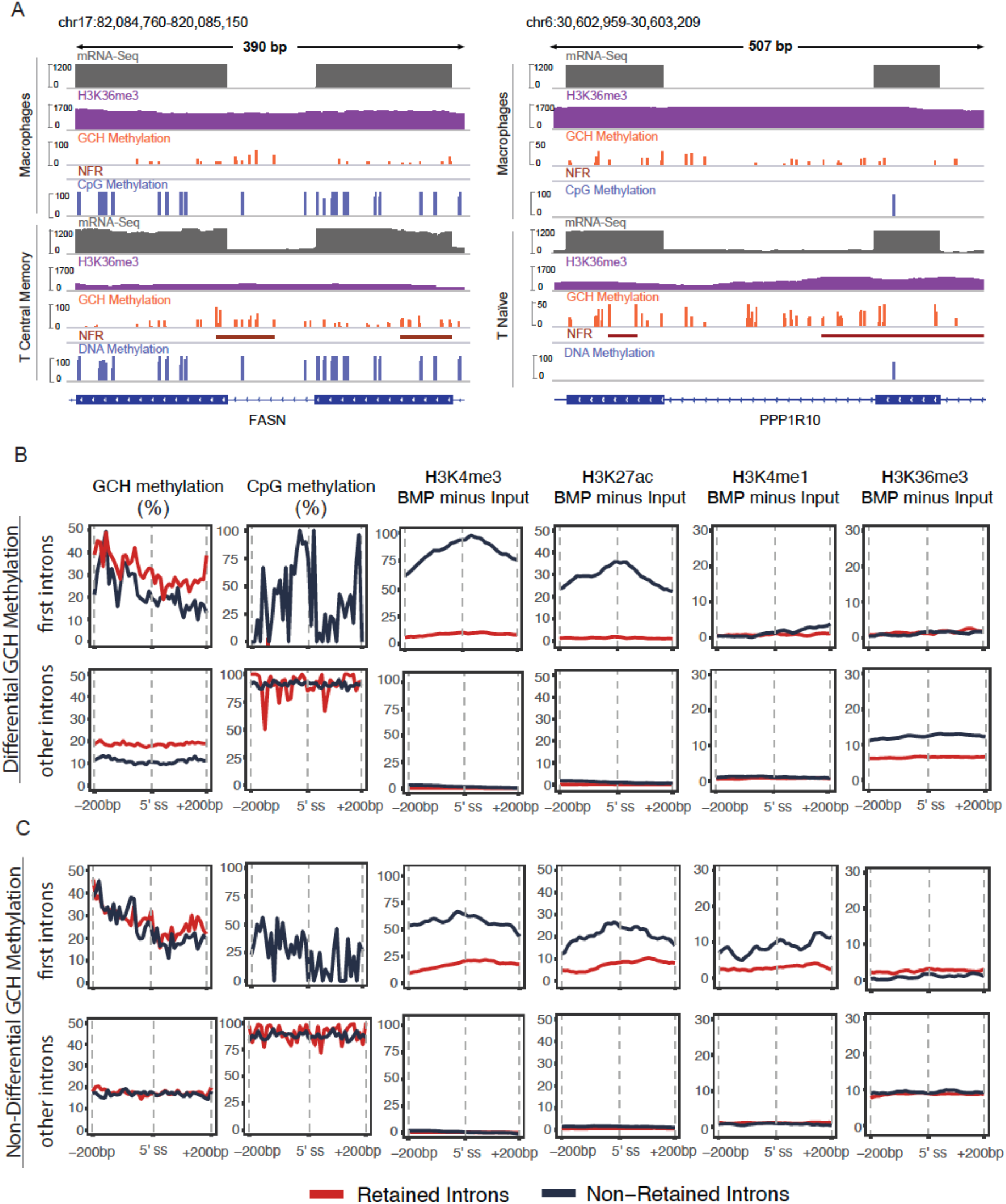
Interplay between chromatin accessibility, CpG methylation and histone modifications. **(A)** IGV plots of mRNA-seq, H3K36me3 ChIP-seq, NOMe-seq, and WGBS-seq data indicating different levels of GCH methylation between retained and non-retained introns and higher prevalence of NFRs in the regions proximal to IR. **(B)** Line graphs show the average levels of GCH methylation, CpG methylation, and the difference between ChIP-seq H3K4me3, H3K27ac, H3K4me1, and H3K36me3 signals and ChIP-Seq Input, normalised to the Bins Per Million (BPM), in retained (red) and non-retained (blue) introns associated with chromatin status. The first row shows epigenetic signals at the 5’ splice site of first introns (close to the promoter region) and the second row represents all other introns. **(C)** The same analysis performed in **(B)** is repeated for introns where the chromatin status remains the same, i.e. non-differential GCH methylation.

Based on the observed patterns, we hypothesise that there is an association between chromatin dynamics and IR: chromatin is more likely to be in a permissive state (high GCH methylation) in the vicinity of retained introns and more compact (low GCH methylation) around constitutively spliced introns. Indeed, we observed that chromatin becomes more accessible as introns become retained (65% of observations). In other cases, the IR status changes without any change to the chromatin state (35% of observations).

Based on the observations concerning chromatin accessibility, we sought to assess the relationship between IR and epigenetic factors in the context of changing chromatin states, i.e. differential GCH methylation (Figure 6B), and stable chromatin status, i.e. non-differential GCH methylation (Figure 6C). In our analysis, we separated first introns from other introns to detach epigenetic signals associated with gene promoters.

The patterns of CpG methylation, H3K27ac, H3K4me3 and H3K4me1 levels in retained and non-retained introns were similar in both chromatin modes (dynamic and stable). First non-retained introns displayed enrichment for histone marks and reduced CpG methylation levels, while first retained introns had negligible levels of histone marks and were marked by the absence of CpG methylation (Figure 6B and 6C, top rows). In contrast, the above-mentioned histone marks were silenced in the internal introns irrespective of the IR status, while the H3K36me3 signal increased. Interestingly, H3K36me3 levels were reduced in retained introns associated with dynamic chromatin (Figure 6B, 2^nd^ row, far right), while they remained similar in retained- and non-retained introns associated with stable chromatin (Figure 6C, 2^nd^ row, far right).

A most interesting result of this analysis was that there are no differences in epigenetic marks between internal retained and non-retained introns when a stable chromatin state is maintained (Figure 6C, bottom row). This suggests that there must be unknown factors that are independent of chromatin accessibility responsible for modulating IR. Thus, further investigations are required to identify additional factors that impact on IR in haematopoietic cells.

## Discussion

In this study, we have employed a machine-learning approach to determine regulators of IR in primary hematopoietic cells. For the first time we provide integrated matched transcriptomic, nucleosome occupancy, CpG methylation, and 6 histone modification profiles from 5 primary human cell types representing 2 independent systems of haematopoietic cell differentiation. Previous studies have described features that are associated with retained introns, including a higher intronic GC content, shorter intron lengths, weaker 5’ and 3’ splice site strengths, and some epigenetic marks (Braunschweig et al., 2014; Schmitz et al., 2017; Wong et al., 2017a). However, these studies have focused on single or paired omics layers only and often used individual cell lines for their analyses.

We applied supervised machine learning using EN and conditional RF algorithms. Unlike deep learning methods, that are very capable of identifying complex relationship patterns but do not provide tools to determine how exactly an outcome was determined (Rauschert et al., 2020), these multivariate models allows the identification of features that contribute most to the outcome of interest (IR). Such modelling strategy is “data-independent” and can be applied to other forms of alternative splicing as well. For example, RF has been used to study the importance of chromatin modifications in the interaction between topologically associated domains (Dixon et al., 2015) and EN was used to model prognostic alternative splicing signatures in breast cancer (Wang et al., 2020).

Previous studies have mostly focussed on investigating the functional links between chromatin organisation and gene expression regulation and found that nucleosome free regions at a transcription start site are strongly associated with transcription initiation (Radman-Livaja and Rando, 2010). Nucleosomes were also reported to be preferentially positioned in exons to facilitate their identification among flanking introns by the splicing machinery (Schwartz et al., 2009; Tilgner et al., 2009). However, it is important to note that these findings were made using the micrococcal nuclease digestion with deep sequencing (MNase-seq) protocol, which is more susceptible to GC content bias. Kelly et al. (Kelly et al., 2012) showed that nucleosome enrichment in exons vs. introns was not observed in NOMe-seq data, which they attributed to the technical differences between the two experimental approaches. NOMe-seq data includes the percentage of methylated reads at a given position as opposed to the count of mapped reads in MNase-seq data. Similarly, our NOMe-seq based analysis of chromatin accessibility, quantified by GCH methylation, did not reveal a specific preference for nucleosomes to be positioned in exons rather than introns.

Our study did reveal the regions of clear GCH enrichment clusters either upstream, downstream or directly at the splice sites of retained introns in contrast to non-retained introns. High GCH methylation levels, like those observed in retained introns, are indicative of nucleosome free regions or NFRs, regions of possible nucleosome eviction that are characterised by a high density of methylated GCH sites and unmethylated CpG dinucleotides (Nordström et al., 2019). Interestingly, You et al. showed that a loss of nucleosome depleted regions accompanied by nucleosome occupancy precedes changes in endogenous CpG methylation in OCT4 and NANOG genes in embryonic carcinoma cell line NCCIT (You et al., 2011). Formation of an NFR upstream from the 5’ exon/intron boundary led to DNA hypomethylation and the depletion of H3K36me3 in SETD2 deficient tumours (Simon et al., 2014). It is therefore reasonable to conclude that alteration of the epigenetic landscape attributed to IR initially starts with changes in nucleosome architecture and subsequent transcriptome rewiring.

Apart from signalling a nucleosome eviction, high levels of GCH methylation potentially mark regions with longer internucleosomal spacing, also known as DNA linker regions. A study in estimating nucleosome phasing in single cell found great agreement between average linker length measured with scNOMe-seq data and the phase estimates derived from MNase-seq (Pott, 2017). Linker length ranges between ~20-90 bp and varies among different species, tissues, and even fluctuates within a single cellular genome (Szerlong and Hansen, 2011). Nucleosome phasing has been linked to alternative splicing before, where RNA Pol II elongation rates increase upon histone depletion and pre-mRNA splicing is delayed (Jimeno-González et al., 2015). Previous studies identified nucleosomes as physical barriers to efficient transcription elongation *in vitro*, however *in vivo* they are efficiently removed from transcribed chromatin (Saldi et al., 2016). Pol II were also found to be involved in maintaining nucleosome phasing in the transcribed region, where longer Pol II dwell times, associated with slow transcription, allowed for remodelling of H3K36me3 profiles (Fong et al., 2017).

In regions further downstream of transcription start sites, nucleosome positioning becomes less stable (Radman-Livaja and Rando, 2010) and linker region lengths become nonuniform. We therefore propose that the differences in DNA methylation and H3K36me3 signal observed over internal introns reflect the underlying changes in nucleosome organisation, that in turn propagates IR (Figure 7). In the presence of IR, transcription rates are faster over more spaced out nucleosomes that does not allow sufficient time for a “writer” to deposit H3K36me3 in the splicing region (Fong et al., 2017). CpG sites in the DNA linker regions are usually unmethylated (Pott, 2017) and therefore may explain the reduced DNA methylated levels associated with IR (Wong et al., 2017b).

**Figure 7.**
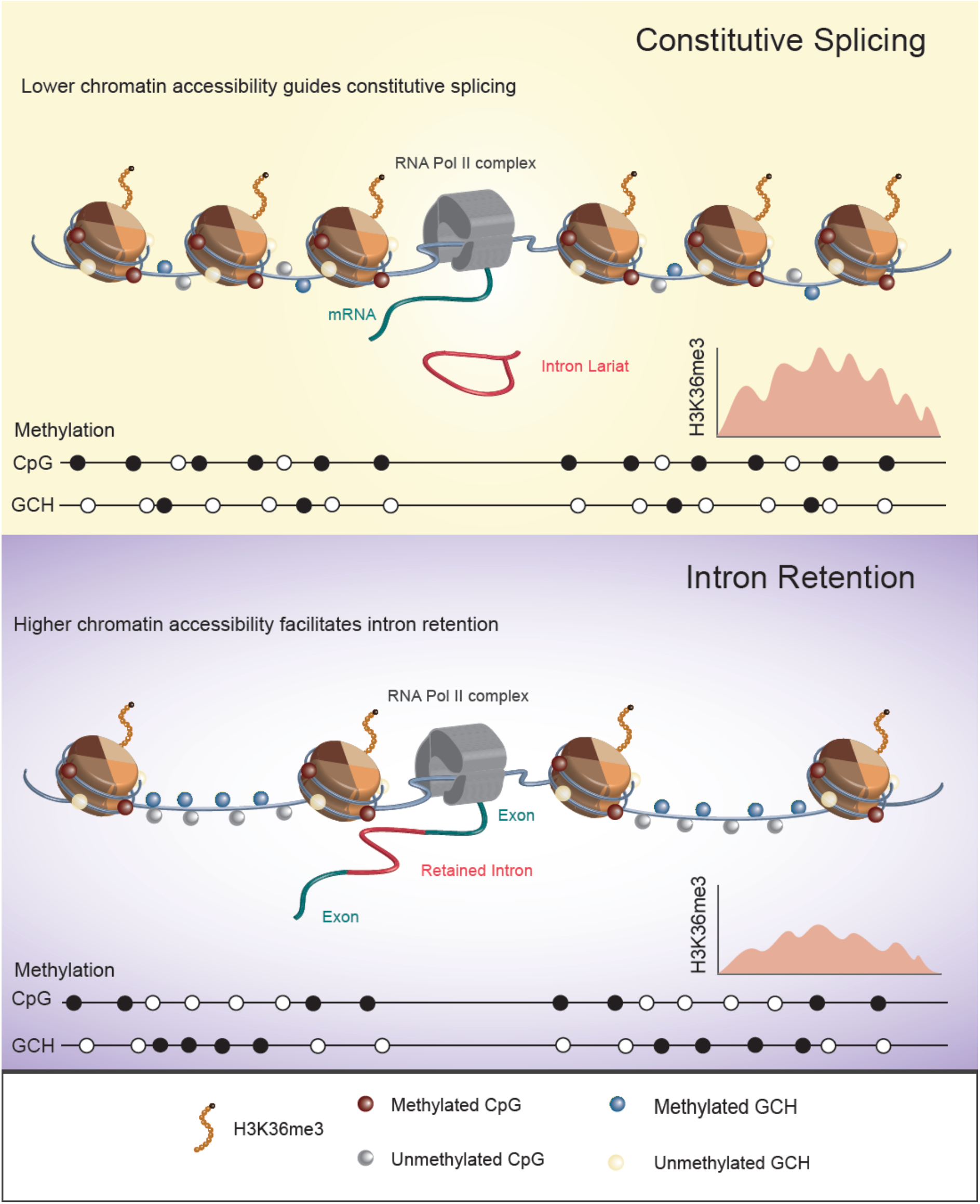
Proposed role of chromatin accessibility in IR regulation. More dense positioning of nucleosomes slows down RNA Pol II elongation rate, allowing sufficient time for a histone modification (in this case, H3K36me3). Methylated CpG dinucleotides and unmethylated GCH sites over the nucleosome core explain higher CpG methylation levels and lower GCH methylation levels in constitutively spliced introns.

In the proximity of transcription start sites, strong histone modification levels (like we observed for H3K4me3 and H3K27ac) indicate a well-positioned nucleosome (Andersson et al., 2009), while reduced histone modification levels, particularly reduced H3K4me3, are associated with transcription factor (TF) binding (Wu et al., 2015). TF binding sites can undergo nucleosome remodelling (Ballaré et al., 2013) in the form of nucleosome shifts or nucleosome eviction and the formation of an NFR with associated changes to RNA polymerase II elongation rates. We propose that IR in first introns might be a biproduct of functional histone modifications and nucleosome remodelling for the purpose of TF recruitment in the regions proximal to transcription start sites.

In summary, our results provide a major conceptual advance in our understanding of alternative splicing regulation. We found an unanticipated strong contribution of chromatin organisation in IR modulation where nucleosomes position upstream or downstream of retained introns (determined by the length of linker regions and NFRs) to facilitate an acceleration of RNA Pol II elongation and increased IR. Furthermore, the models generated in this study can be adapted to study epigenetic gene expression and alternative splicing regulation in other cell systems, other species, in health or disease, and further our understanding of these essential biological mechanisms.

## Acknowledgements

We thank Benedikt Brors and Roland Eils from DKFZ Heidelberg and Alf Hamann from DRFZ Berlin, Wie Chen, Nikolaus Rajewsky and Sascha Sauer from MDC Berlin, Ho-Ryun Chung and Martin Vingron from MPI-MG Berlin, Thomas Jenuwein, Thomas Manke and Andrew Pospisilik from MPI-IE Freiburg, Philip Rosenstiel and Stefan Schreiber from CAU Kiel, Jan G. Hengstler from IfADo Dortmund, Thomas Lengauer from MPI-INF Saarbrücken, Bernhard Horsthemke from Universität Duisburg-Essen, Alexandra Kiemer from Universität des Saarlandes Saarbrücken, Thomas Pap from WWU Münster and Gerd Schmitz from Universität Regensburg who were involved in the work with biological samples, sequencing and generation of WGBS, NOMe-Seq, ChIP-Seq and RNA-Seq data for the DEEP Consortium.

This work was supported by the National Health and Medical Research Council (Investigator Grant #1177305 to J.E.J.R., Project #1080530 to J.E.J.R., Project #1128175 and #1129901 to J.E.J.R. and J.J.-L.W., #1126306 to J.J.-L.W.; the NSW Genomics Collaborative Grant (J.E.J.R. and J.J.-L.W.); Cure the Future (J.E.J.R.), and an anonymous foundation (J.E.J.R.). U.S. and J.J.-L.W. hold Fellowships from the Cancer Institute of New South Wales. U.S. also received support from the Australian Academy of Science in form of an Australia-India Early and Mid-Career Fellowship. This research was funded by the Cancer Council NSW Project Grants (RG11-11 and RG20-12) to J.E.J.R. and U.S. K.V.J.N. and J.W. were supported by the German Epigenome Program (DEEP) funded by the Ministry of Education and Research in Germany (BMBF 01KU1216).

The authors acknowledge the technical assistance provided by the Sydney Informatics Hub, a Core Research Facility of the University of Sydney.

## Author Contributions

J.E.J.R., J.J.-L.W. and U.S. designed the study and supervised the project, V.P. and R.S. performed bioinformatic analyses, V.P. performed statistical analysis and data modelling, N.J.A. advised on statistical methodology, DEEP Consortium provided sequencing data, J.W. designed and coordinated sequencing experiments, K.J.V.N. data management, V.P. and U.S. wrote the manuscript. All authors have read and agreed to the published version of the manuscript.

## STAR Methods

### Quantification and statistical analysis

To investigate how IR is regulated in primary immune cells, we integrated epigenomics and transcriptomics data from the German Epigenome Program (DEEP). Primary monocytes, monocyte-derived macrophages, and primary T-cells (naïve, central memory, effector memory) were retrieved from 2 healthy donors. Cell isolation, differentiation, DNA/RNA extraction and library preparation for mRNA-Seq, WGBS, NOMe- and ChIP-Seq experiments are described in detail in these articles (Durek et al., 2016; Wallner et al., 2016).

#### mRNA-Seq data processing and identification of IR events

RNA-Seq reads (FASTQ format) of each technical replicate were tested for quality using FastQC v.0.11.5 (github.com/s-andrews/FastQC). Further processing, including adaptor trimming, was performed within the IRFinder algorithm for IR quantification (Middleton et al., 2017). Sequencing reads were mapped to the human reference genome (GRCh38) using STAR v2.7 with default parameters (Dobin et al., 2013). IR-ratios, a quantitative measure of IR levels, were determined as:

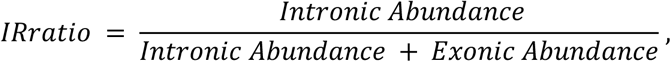

where the Intronic Abundance is defined as the trimmed mean of the reads that map to an intron, after having excluded features that overlap the intron, with the highest and lowest 30% of values being excluded. Exonic Abundance is defined as the number of reads that map across an exon-exon junction. Library size normalisation (between-sample normalisation) was not required as the ratio between intronic and exonic abundance is determined from within the same transcriptome (Middleton et al., 2017).

Introns that were present in at least 10% of a gene’s mature mRNA transcripts (IR_ratio_ ≥ 0.1) with an overall intron depth ≥ 10 were considered retained. Non-retained introns were defined as those with an IR_ratio_ ≤ 0.01 and intron depth < 10.

We used Cufflinks v2.1.1 (Trapnell et al., 2010) to estimate gene abundance in fragments per kilobase per million (FPKM). Only introns from host genes with FPKM ≥ 1 were selected for the downstream analyses.

#### WGBS data processing

Raw WGBS FASTQ files were assessed for quality using FastQC v.0.11.5 (github.com/s-andrews/FastQC). Standard Illumina adaptors used for the library preparation were trimmed using cutadapt v.1.10 (Martin, 2011) with a quality cutoff of 20 base pairs (bp) and minimum read length of 30 bp. Trimmed reads were mapped to the GRCh38 reference genome, duplicate reads removed, and methylation calling performed using Bismark v.0.19.0 (Krueger and Andrews, 2011). Only CpG sites with a coverage of more than 5 reads were retained for further analysis.

#### ChIP-Seq data processing

ChIP-Seq data for six histone modifications (H2K27ac, H3K27me3, H3K36me3, H3K4me1, H3K4me3, H3K9me3) were aligned to the human GRCh38 reference genome using STAR v2.7 (Dobin et al., 2013). Duplicate reads were removed using Picard v.2.18.4 (broadinstitute.github.io/picard/) and further processed using MACS2 v.2.2.6 (Zhang et al., 2008) to identify histone modification peaks, with default parameters and q-value cut-off of 0.01. All histone modifications were processed in the “narrow peak” mode in order to extract peak summit coordinates. For visualisation in IGV (Robinson et al., 2012), we generated coverage tracks using `bamCoverage` from deepTools2 (Ramirez et al., 2016) with the following parameters `--binSize 1 --normalizeUsing BPM --effectiveGenomeSize 2913022398 --extendReads 200`. For HM line plots, we substracted ChiP-Seq Input from a respective HM ChiP-seq read counts and normalised based on Bins Per Million (BPM) mapped reads using bamCompare and parameters `--binSize 1 --scaleFactorsMethod readCount --effectiveGenomeSize 2913022398 --operation subtract --normalizeUsing BPM`.

#### NOMe-Seq data processing

Raw FASTQ files were assessed for quality using FastQC v.0.11.5 (github.com/s-andrews/FastQC). Reads were mapped to the GRCh38 reference genome, duplicate reads removed, and methylation calling performed using Bismark v.0.19.0 (Krueger and Andrews, 2011). GCH methylation information was extracted with the `coverage2cytosine` utility with `--nome` parameter.

NFRs were predicted using gNOMePeaks tool (Nordström et al., 2019) with default parameters, which include 4,000 bp up- and downstream from each peak for background signal calculation and the maximum distance between GpC sites of 150 bp. We used the same algorithms to predict nucleosome positioning by substituting GCH methylation, as required input, with GCH occupancy (1 – *GCH methylation*) and reducing the background region to 1,000 bp up- and downstream from each peak and the distance between GCH sites to 20bp.

#### Feature selection

Model features were associated with three genomic regions around retained and non-retained introns: (i) +/- 100 bp from the 5’splice site, (ii) +/- 100 bp from the 3’splice site, and (iii) +/- 100bp from the middle of an intron, each region being 200 bp long. GC content was extracted using bedtools v.2.26.0 (Quinlan and Hall, 2010) `nuc` command. For splice site strength calculations, we used MaxEntScan (Yeo and Burge, 2004). CpG density values was obtained using Repitools (Statham et al., 2010). The percent spliced in (PSI) index of flanking exons was calculated as described in (Schafer et al., 2015). Exons with PSI ≥ 0.9 were considered as included.

To generate epigenetic features, we overlapped three regions of interest with the pre-processed epigenetic data. NFR regions were defined as regions greater than 40bp in length with p-value ≤ 0.05 (Fisher test comparing CpG methylation in the NFR to the surrounding background). Presence or absence of an NFR was dichotomised as “yes” −1 and “no” −0. Information about nucleosome location was included into the model in the similar manner (nucleosomes were defined as regions greater than 140bp in length with p-value ≤ 0.05).

The relationship between histone modification and IR was included into the model through the presence or absence of an overlap with a histone signal region. It was categorised as 0 – no overlap, 1 – overlap with a region of HM signal, 2 – overlap with a region of strong signal (strong signal = mean (HM pile-up) + sd (HM pile-up)). The full list of features is presented in Table S1.

### Elastic Net and Conditional Random Forest Modelling

To identify features important for IR, we constructed a binary classification model using the EN algorithm. We approached the problem in a naïve manner, i.e. we did not impose any prior assumptions about the factors that might potentially play a role and therefore an equal penalty factor was applied to all features. EN classification was performed in the *caret* R package (Kuhn, 2008) using *glmnet* method (Friedman J, 2010) for a binary outcome. The group imbalance, due to the different number of retained and non-retained introns identified suitable for modelling, was handled by down-sampling, using *downSample* command. Parameter *λ*, determining the overall size of the regularization penalty, was optimised by 10-fold cross validation procedure. Features were ranked based on the absolute values of the model coefficients.

We repeated this *in-silico* analysis to validate our results using an independent machine learning algorithm, cRF. In cRF, unlike standard RF where the first split variable is randomly selected, an association test between the outcome and the model predictors is performed first. The ranked p-values are then used to identify the covariate with the strongest association to the outcome, which is later used for the first binary split at cutpoint *c* for a continuous covariate or at category *C* for a categorical covariate. cRF classification was also performed in *caret* using *cforest* method as implemented in the *party* R package (Strobl C, 2008). The cRF model provides an unbiased measure of variable importance, which we used to rank the most important features for IR prediction.

To avoid overfitting, we ranked the features’ importance using both EN and cRF techniques (Ding et al., 2018). Moreover, our findings were validated across different blood cell lineages from different humans.

### Statistical Analysis

All statistical analyses were performed in R v.4.0. For the identification of differentially retained introns we used the Audic and Claverie Test (Audic and Claverie, 1997). P-values ≤ 0.05 were considered significant. Clustering was performed using unsupervised hierarchical clustering with complete linkage.

### Data and Software Availability

Sequencing data are deposited at the European Genome-Phenome Archive under the accession numbers EGAS00001001595 and EGAS00001001624. Access is subject to an application process as per the EGA requirements. R scripts developed for this study are available at https://github.com/combiomed/IR_code. Processed sequencing data used to train the models was deposited at Mendeley Data: http://dx.doi.org/10.17632/b6crxbxbk2.1.

