## Supplemental Materials for "Chromatin accessibility determines intron retention in a cell type-specific manner"

### Supplementary Tables

**Table S1 Experiments included in the study (separate Excel File)**

**Table S2 Epigenetic signals intersecting with 5´ splice site regions (+/- 100 bp).**

| | | **Monoc** | | **Macro** | | **Naïve T** | | **T-CM** | | **T-EM** | |
|---|---|---|---|---|---|---|---|---|---|---|---|
| | | Ret | Non-Ret | Ret | Non-Ret | Ret | Non-Ret | Ret | Non-Ret | Ret | Non-Ret |
| **H3K9me3** | Strong Signal | 0 | 22 | 2 | 41 | 0 | 0 | 0 | 0 | 0 | 0 |
| | Signal | 6 | 107 | 5 | 84 | 3 | 11 | 1 | 1 | 13 | 36 |
| | No Signal | 1,633 | 28,267 | 812 | 30,845 | 4,816 | 20,671 | 4,944 | 18,928 | 5,634 | 18,227 |
| **H3K27me3** | Strong Signal | 7 | 21 | 2 | 11 | 0 | 0 | 0 | 0 | 0 | 0 |
| | Signal | 9 | 108 | 5 | 92 | 0 | 2 | 1 | 4 | 0 | 1 |
| | No Signal | 1,623 | 28,267 | 812 | 30,867 | 4,819 | 20,680 | 4,944 | 18925 | 5,647 | 18,262 |
| **H3K27ac** | Strong Signal | 46 | 596 | 23 | 1,004 | 8 | 9 | 0 | 0 | 22 | 27 |
| | Signal | 79 | 964 | 41 | 1,094 | 50 | 201 | 36 | 66 | 150 | 291 |
| | No Signal | 1,514 | 26,836 | 755 | 28,872 | 4,761 | 20,472 | 4,909 | 18,863 | 5,475 | 17,945 |
| **H3K36me3** | Strong Signal | 550 | 3,235 | 217 | 4,983 | 502 | 660 | 1,379 | 1,005 | 814 | 577 |
| | Signal | 380 | 10,150 | 229 | 11,621 | 1,411 | 4,588 | 1,459 | 5,260 | 849 | 2,754 |
| | No Signal | 709 | 15,011 | 373 | 14,366 | 2,906 | 15,434 | 2,107 | 12,664 | 3,984 | 14,932 |
| **H3K4me1** | Strong Signal | 52 | 187 | 31 | 250 | 102 | 31 | 91 | 26 | 151 | 43 |
| | Signal | 131 | 1,009 | 87 | 1,428 | 301 | 333 | 375 | 274 | 438 | 361 |
| | No Signal | 1,456 | 27,200 | 701 | 29,292 | 4,416 | 20,318 | 4,479 | 18,629 | 5,058 | 17,859 |
| **H3K4me3** | Strong Signal | 40 | 1,099 | 19 | 1,244 | 71 | 297 | 85 | 365 | 89 | 314 |
| | Signal | 61 | 301 | 26 | 437 | 133 | 526 | 168 | 449 | 185 | 400 |
| | No Signal | 1,538 | 26,996 | 774 | 29,289 | 4,615 | 19,859 | 4,692 | 18,115 | 5,373 | 17,549 |
| **NFR** | Peak | 202 | 7,841 | 90 | 7,057 | 1,502 | 5,366 | 1,528 | 4,204 | 2,052 | 5,255 |
| | No Peak | 1,437 | 20,555 | 729 | 23,913 | 3,317 | 15,316 | 3,417 | 14,725 | 3,595 | 13,008 |

**Table S3 Features used for modelling IR.**

| Abbreviation | Feature Description |
|---|---|
| **Intrinsic Features (mRNA-Seq)** | |
| Length | Intron length (continuous, bp) |
| Length_5ssexon | Upstream exon length (continuous, bp) |
| Length_3ssexon | Downstream exon length (continuous, bp) |
| GCcont_int | Intron GC content (proportional, 0-1) |
| GCcont_5ssexon | Upstream exon GC content (proportional, 0-1) |
| GCcont_3ssexon | Downstream exon GC content (proportional, 0-1) |
| MaxEntScore5ss | 5′ splice site strength (continuous) |
| MaxEntScore3ss | 3′ splice site strength (continuous) |
| SplicedIn_5ssexon | Upstream exon is spliced in (No – 0, Yes – 1) |
| SplicedIn_3ssexon | Downstream exon is spliced in (No – 0, Yes – 1) |
| **CpG methylation Features (WGB-Seq)** | |
| avemethint_100bp5ss | Average CpG methylation in the region +/- 100 bp from 5′ splice site (proportional, 0-1) |
| avemethint_100bp3ss | Average CpG methylation in the region +/- 100 bp from 3′ splice site (proportional, 0-1) |
| avemethint_100bpmid | Average CpG methylation in the region +/- 100 bp at the middle of an intron (proportional, 0-1) |
| CpGint_100bp5ss | Number of CpG sites in the region +/- 100 bp from 5′ splice site (count) |
| CpGint_100bp3ss | Number of CpG sites in the region +/- 100 bp from 3′ splice site (count) |
| CpGint_100bp5ss | Number of CpG sites in the region +/- 100 bp from the middle of an intron (count) |
| **Nucleosome Occupancy Features (NOMe-Seq)** | |
| gnomePeak_olap_100bp5ss | Overlap with nucleosome peak in the region +/- 100 bp from 5′ splice site (0 – No, 1 – Yes) |
| gnomePeak_olap_100bp3ss | Overlap with nucleosome peak in the region +/- 100 bp from 3′ splice site (0 – No, 1 – Yes) |
| gnomePeak_olap_100bpmid | Overlap with nucleosome peak in the region +/- 100 bp from the middle of an intron (0 – No, 1 – Yes) |
| gnomePeak_olap_NFR_100bp5ss | Overlap with NFR in the region +/- 100 bp from 5′ splice site (0 – No, 1 – Yes) |
| gnomePeak_olap_NFR_100bp3ss | Overlap with NFR in the region +/- 100 bp from 3′ splice site (0 – No, 1 – Yes) |
| gnomePeak_olap_NFR_100bpmid | Overlap with NFR in the region +/- 100 bp from the middle of an intron (0 – No, 1 – Yes) |
| GCHmeth_100bp5ss | Average GCH methylation in the region +/- 100 bp from 5′ splice site (proportional, 0-1) |
| GCHmeth_100bp3ss | Average GCH methylation in the region +/- 100 bp from 3′ splice site (proportional, 0-1) |
| GCHmeth_100bpmid | Average GCH methylation in the region +/- 100 bp from the middle of an intron (proportional, 0-1) |
| GCHoccup_100bp5ss | Average GCH occupancy in the region +/- 100 bp from 5′ splice site (proportional, 0-1) |
| GCHoccup_100bp3ss | Average GCH occupancy in the region +/- 100 bp from 3′ splice site (proportional, 0-1) |
| GCHoccup_100bpmid | Average GCH occupancy in the region +/- 100 bp from the middle of an intron (proportional, 0-1) |

| **Histone Modification Features (ChIP-Seq)** | |
|---|---|
| H3K4me1_olap_100bp5ss | Overlap with H3K4me1 signal in the region +/- 100 bp from 5′ splice site (0 – no signal, 1 – signal, 2 – strong signal) |
| H3K4me3_olap_100bp5ss | Overlap with H3K4me3 signal in the region +/- 100 bp from 5′ splice site (0 – no signal, 1 – signal, 2 – strong signal) |
| H3K9me3_olap_100bp5ss | Overlap with H3K9me3 signal in the region +/- 100 bp from 5′ splice site (0 – no signal, 1 – signal, 2 – strong signal) |
| H3K27ac_olap_100bp5ss | Overlap with H3K27ac signal in the region +/- 100 bp from 5′ splice site (0 – no signal, 1 – signal, 2 – strong signal) |
| H3K27me3_olap_100bp5ss | Overlap with H3K27me3 signal in the region +/- 100 bp from 5′ splice site (0 – no signal, 1 – signal, 2 – strong signal) |
| H3K36me3_olap_100bp5ss | Overlap with H3K36me3 signal in the region +/- 100 bp from 3′ splice site (0 – no signal, 1 – signal, 2 – strong signal) |
| H3K4me1_olap_100bp3ss | Overlap with H3K4me1 signal in the region +/- 100 bp from 3′ splice site (0 – no signal, 1 – signal, 2 – strong signal) |
| H3K4me3_olap_100bp3ss | Overlap with H3K4me3 signal in the region +/- 100 bp from 3′ splice site (0 – no signal, 1 – signal, 2 – strong signal) |
| H3K9me3_olap_100bp3ss | Overlap with H3K9me3 signal in the region +/- 100 bp from 3′ splice site (0 – no signal, 1 – signal, 2 – strong signal) |
| H3K27ac_olap_100bp3ss | Overlap with H3K27ac signal in the region +/- 100 bp from 3′ splice site (0 – no signal, 1 – signal, 2 – strong signal) |
| H3K27me3_olap_100bp3ss | Overlap with H3K27me3 signal in the region +/- 100 bp from 3′ splice site (0 – no signal, 1 – signal, 2 – strong signal) |
| H3K36me3_olap_100bp3ss | Overlap with H3K36me3 signal in the region +/- 100 bp from 3′ splice site (0 – no signal, 1 – signal, 2 – strong signal) |
| H3K4me1_olap_100bpmid | Overlap with H3K4me1 signal in the region +/- 100 bp from the middle of an intron (0 – no signal, 1 – signal, 2 – strong signal) |
| H3K4me3_olap_100bpmid | Overlap with H3K4me3 signal in the region +/- 100 bp from the middle of an intron (0 – no signal, 1 – signal, 2 – strong signal) |
| H3K9me3_olap_100bpmid | Overlap with H3K9me3 signal in the region +/- 100 bp from the middle of an intron (0 – no signal, 1 – signal, 2 – strong signal) |
| H3K27ac_olap_100bpmid | Overlap with H3K27ac signal in the region +/- 100 bp from the middle of an intron (0 – no signal, 1 – signal, 2 – strong signal) |
| H3K27me3_olap_100bpmid | Overlap with H3K27me3 signal in the region +/- 100 bp from the middle of an intron (0 – no signal, 1 – signal, 2 – strong signal) |
| H3K36me3_olap_100bp5ss | Overlap with H3K36me3 signal in the region +/- 100 bp from 5′ splice site (0 – no signal, 1 – signal, 2 – strong signal) |

**Table S4 Model performances when data is trained in one cell type and tested in another.**

Elastic Net Fitted models

| Trained | **Monoc** | | **Macro** | | **Naïve T** | | **T-CM** | | **T-EM** | |
|---|---|---|---|---|---|---|---|---|---|---|
| Tested | AUC | Acc | AUC | Acc | AUC | Acc | AUC | Acc | AUC | Acc |
| **Mono** | 0.923 | 0.890 | 0.917 | 0.922 | 0.861 | 0.774 | 0.930 | 0.908 | 0.891 | 0.908 |
| **Macro** | 0.869 | 0.881 | 0.874 | 0.849 | 0.802 | 0.753 | 0.816 | 0.838 | 0.847 | 0.869 |
| **Naïve T** | 0.891 | 0.905 | 0.900 | 0.911 | 0.947 | 0.950 | 0.939 | 0.947 | 0.944 | 0.949 |
| **T-CM** | 0.907 | 0.913 | 0.922 | 0.923 | 0.933 | 0.868 | 0.954 | 0.958 | 0.946 | 0.949 |
| **T-EM** | 0.898 | 0.899 | 0.922 | 0.923 | 0.935 | 0.869 | 0.942 | 0.946 | 0.948 | 0.951 |

Conditional Random Forest Fitted models

| | **Monoc** | | **Macro** | | **Naïve T** | | **T-CM** | | **T-EM** | |
|---|---|---|---|---|---|---|---|---|---|---|
| | AUC | Acc | AUC | Acc | AUC | Acc | AUC | Acc | AUC | Acc |
| **Mono** | 0.942 | 0.912 | 0.907 | 0.918 | 0.647 | 0.576 | 0.895 | 0.904 | 0.894 | 0.910 |
| **Macro** | 0.870 | 0.879 | 0.905 | 0.867 | 0.787 | 0.778 | 0.816 | 0.844 | 0.842 | 0.874 |
| **Naïve T** | 0.900 | 0.903 | 0.894 | 0.901 | 0.971 | 0.946 | 0.944 | 0.945 | 0.945 | 0.951 |
| **T-CM** | 0.916 | 0.918 | 0.917 | 0.925 | 0.914 | 0.839 | 0.976 | 0.950 | 0.959 | 0.960 |
| **T-EM** | 0.916 | 0.914 | 0.921 | 0.922 | 0.939 | 0.931 | 0.943 | 0.946 | 0.975 | 0.946 |

#### Supplementary Figures

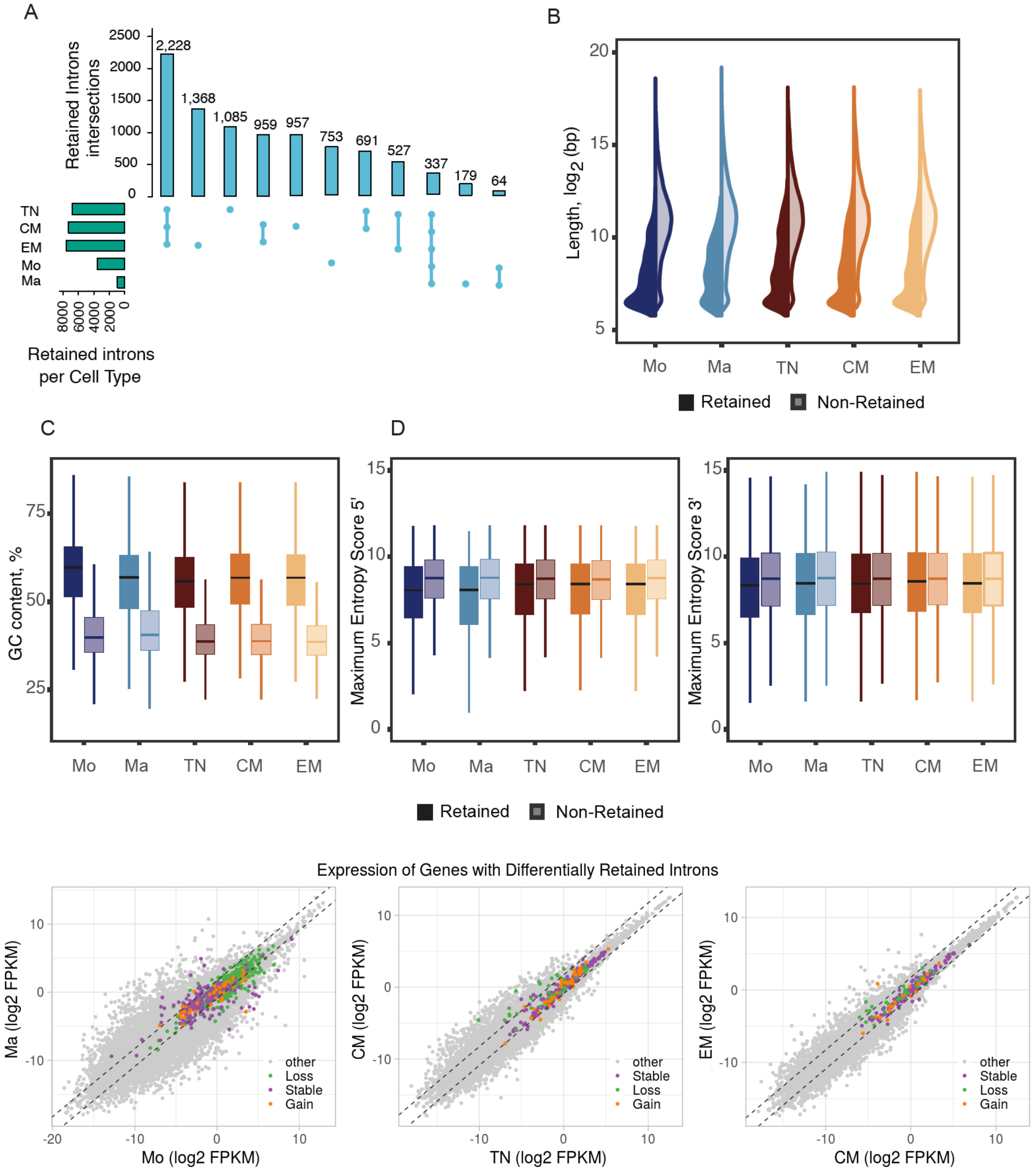


**Figure S1 (A)** UpSet plot of IR event intersections between five cell types: monocytes (Mo), macrophages (Ma), T Naïve (TN), T Central memory (CM) and T Effector Memory (EM). **(B)** Violin plot comparing length distributions between retained and non-retained introns. **(C)** Box plot of average GC content. **(D)** Average 5´ and 3´ splice site strength comparison, measured by Maximum Entropy Score. **(E)** Expression profiles of the genes harbouring introns that are differentially retained during monocyte-to-macrophage differentiation or T cell maturation. Purple – host genes of introns with stable IR, green – host genes of introns with IR loss, orange – host genes of introns with IR gain.

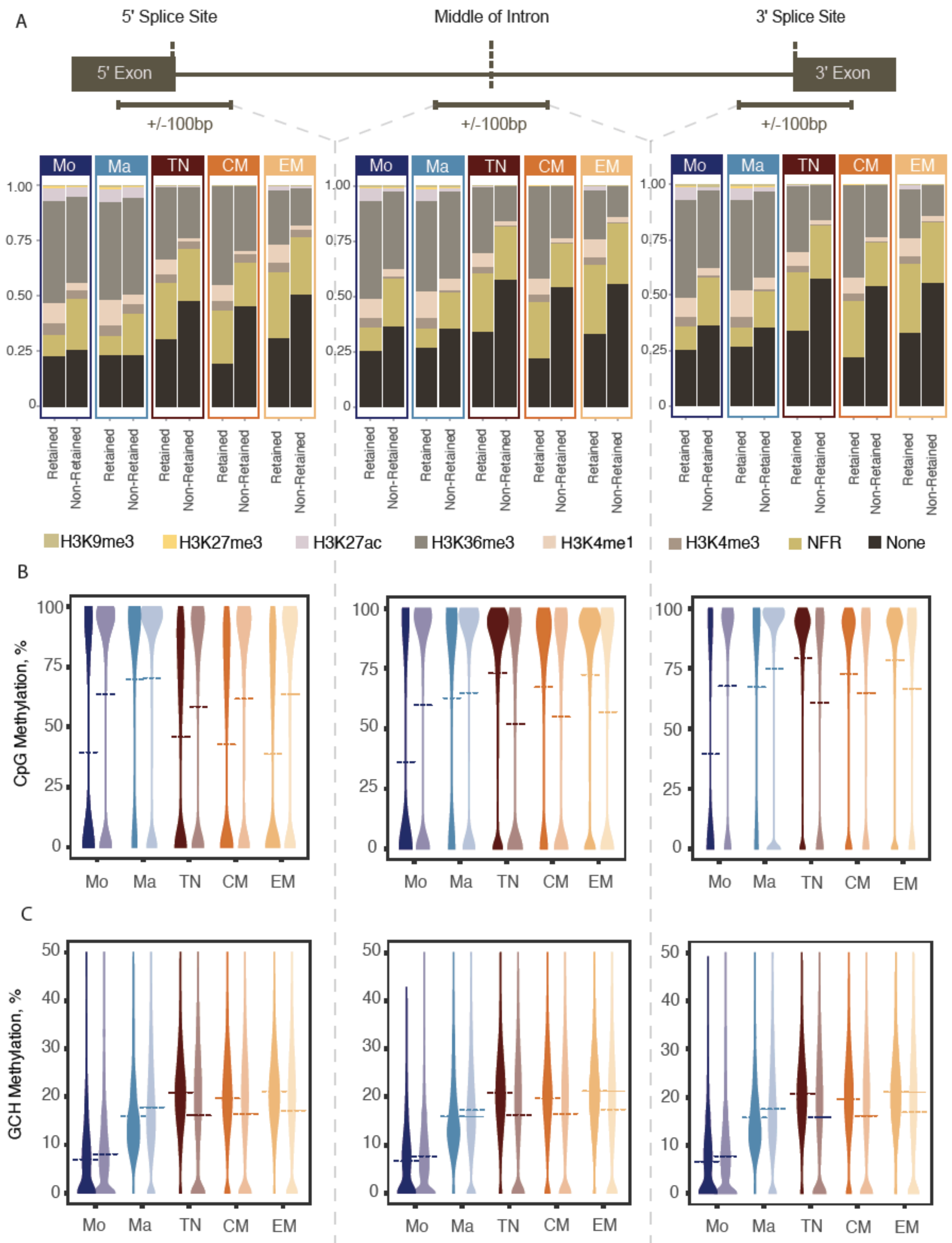


**Figure S2 (A)** Relative count of the HM and NFR peaks overlapping with the 5´ splice site region (left), middle (centre), and 3´ splice site region (right) of retained and non-retained introns. **(B)** CpG methylation distribution patterns at the 5´ splice site (left), middle (centre), and 3´ splice site (right) of retained and non-retained introns. **(C)** GCH methylation distribution patterns at the 5´ splice site (left), middle (centre), and 3´ splice site (right) of retained and non-retained introns.

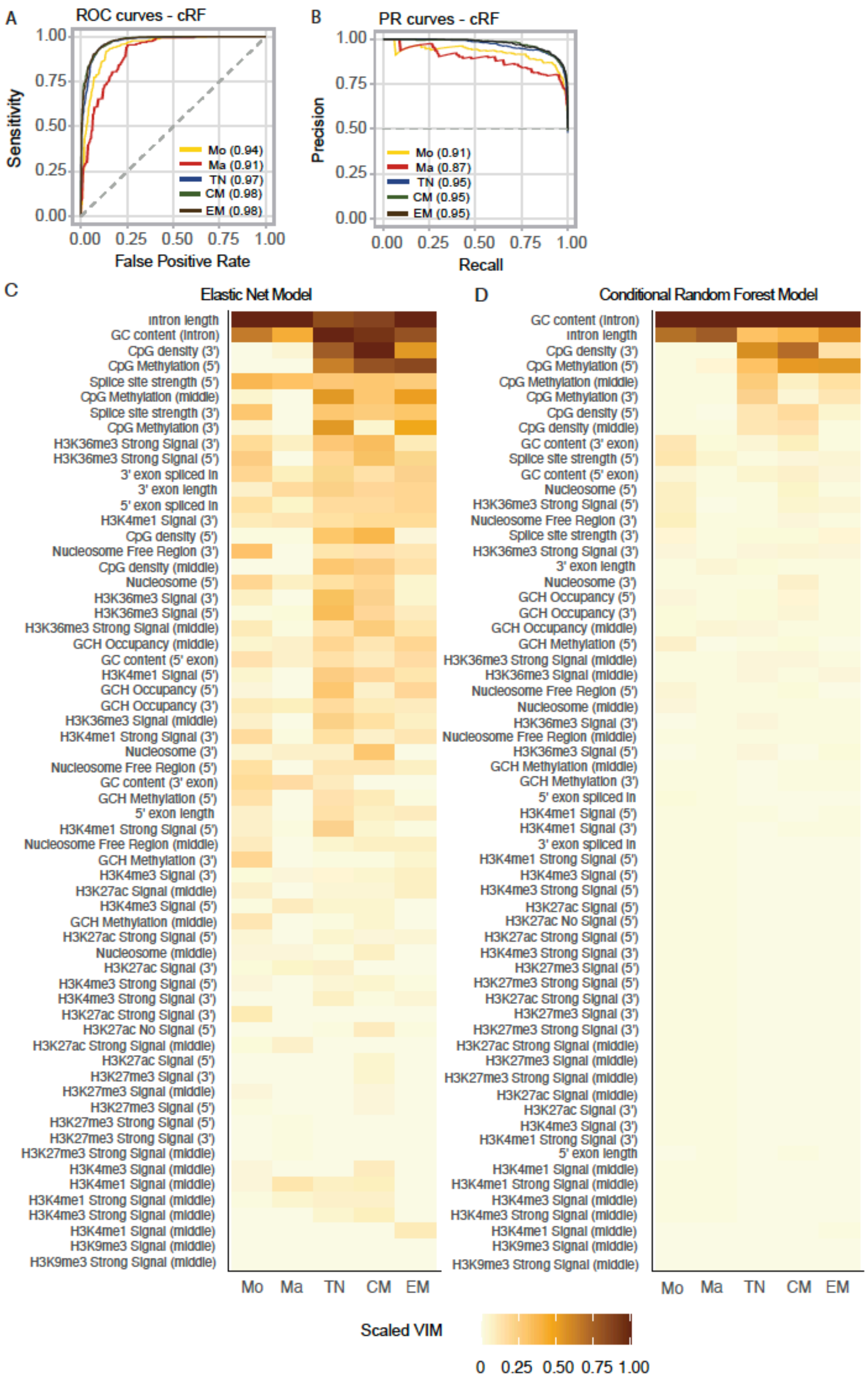

**Figure S3 (A)** ROC curves and **(B)** PR curves illustrating the performance of the cRF classifier in five cell types. Heatmap of scaled VIM values for EN models (C) and cRF models (D).

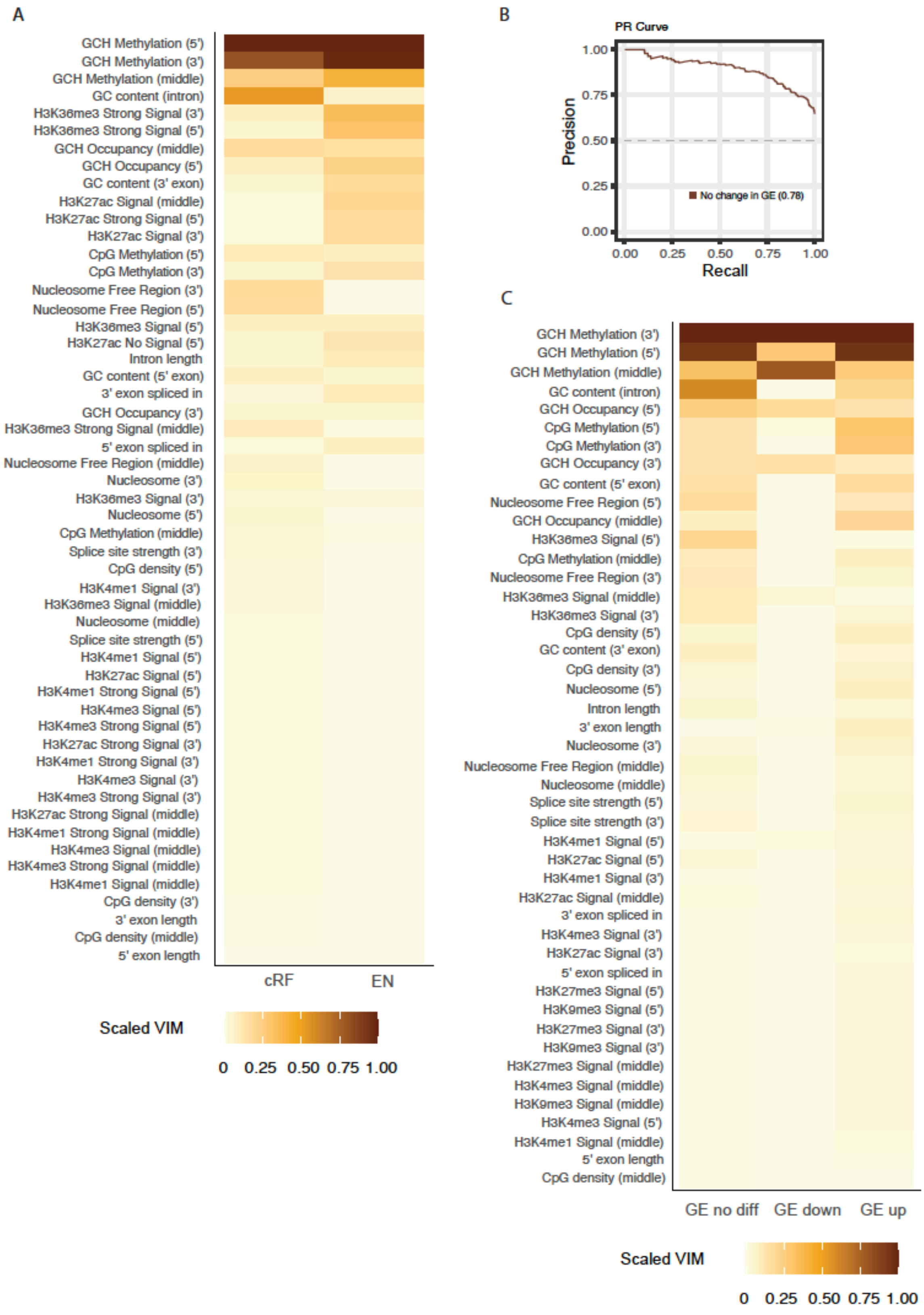


**Figure S4 (A)** Heatmap of scaled VIM values for cRF and EN models run on the dynamic introns subset. **(B)** PR curve for cRF model on dynamic introns with non-differentially expressed host genes. **(C)** Heatmap of scaled VIM values for cRF models run on three groups of dynamic introns: within non-differentially expressed genes, within downregulated genes, and within upregulated host genes.

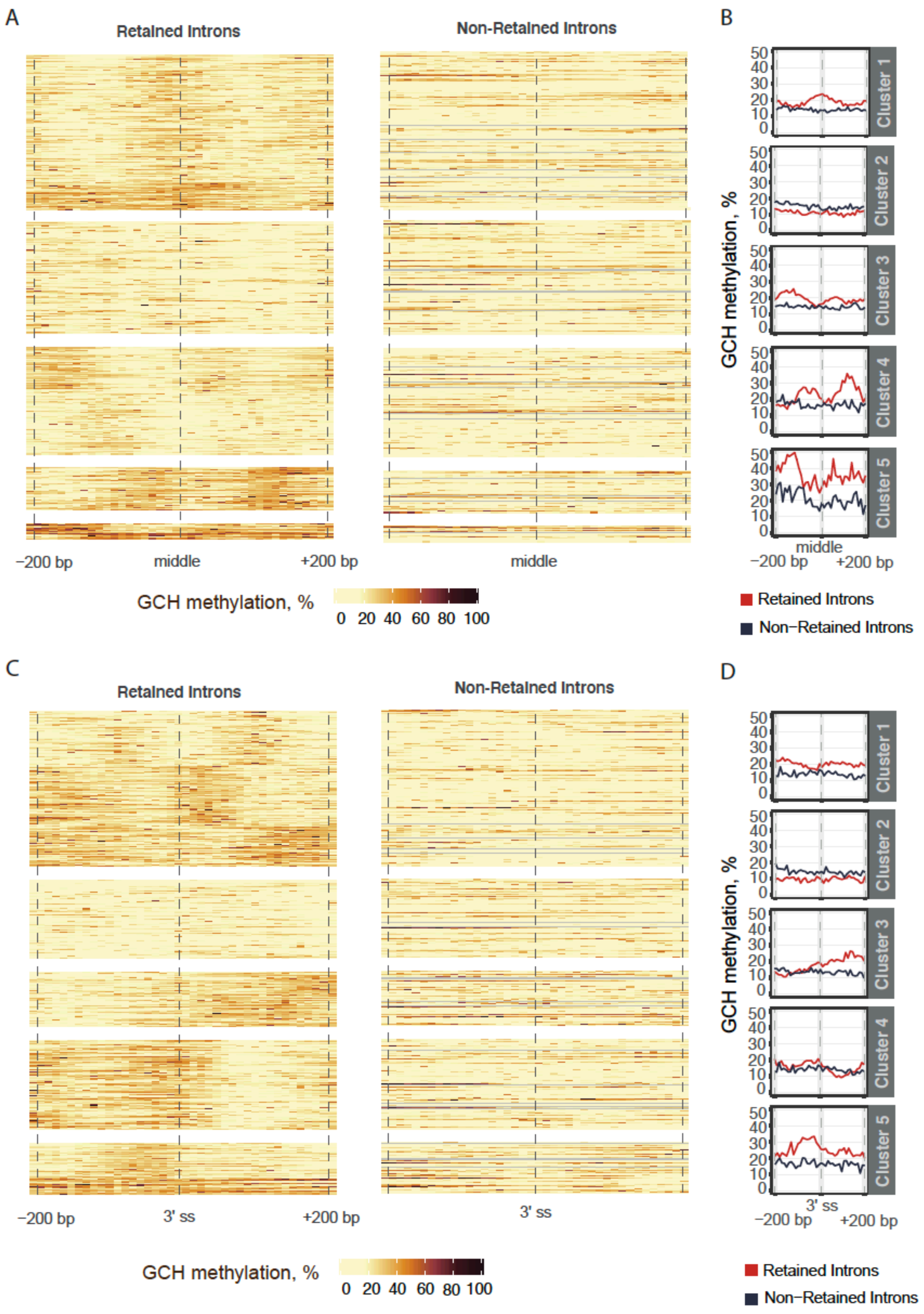


**Figure S5** Clusters of GCH methylation patterns around **(A)** the middle (+/- 200 bp) and **(C)** around the 3' ss (+/- 200 bp) of retained and non-retained introns. Line plots showing average GCH methylation values (i.e., chromatin accessibility) in retained vs non-retained introns across 5 clusters in (**B)** the middle and **(D)** at the 3' ss.
